# Mouse models for V103I and I251L gain of function variants of the human MC4R display reduced adiposity and are not protected from a hypercaloric diet

**DOI:** 10.1101/2020.08.25.266866

**Authors:** Daniela Rojo, Clara McCarthy, Jesica Raingo, Marcelo Rubinstein

## Abstract

**Objective:** The melanocortin 4 receptor (MC4R) is a G protein-coupled receptor that plays major roles in the central control of energy balance. Loss-of-function mutations of MC4R constitute the most common monogenic cause of early-onset extreme obesity in humans, whereas gain-of-function mutations appear to be protective. In particular, two relatively frequent alleles carrying the non-synonymous coding mutations V103I or I251L have been associated with lower risks of obesity and type-2 diabetes. Although V103I and I251L MC4Rs showed more efficient signaling in transfected cells, their specific effects in live animals remain unexplored. Here, we investigated whether the introduction of V103I and I251L mutations into the mouse MC4R leads to a lean phenotype and provides protection against an obesogenic diet.

**Methods:** Using CRISPR/Cas9, we generated two novel strains of mice carrying single nucleotide mutations into the mouse *Mc4r* which are identical to those present in V103I and I251L MCR4 human alleles, and studied their phenotypic outcomes in mice fed with normal chow or a high-fat diet. In particular, we measured body weight progression, food intake and adiposity. In addition, we analyzed glucose homeostasis through glucose and insulin tolerance tests.

**Results:** We found that homozygous V103I females displayed shorter longitudinal length and decreased abdominal white fat, whereas homozygous I251L females were also shorter and leaner due to decreased weight in all white fat pads examined. Homozygous *Mc4r*^*V103I/V103I*^ and *Mc4r*^*I251L/I251L*^ mice of both sexes showed improved glucose homeostasis when challenged in a glucose tolerance test, whereas *Mc4r*^*I251L/I251L*^ females showed improved responses to insulin. Despite being leaner and metabolically more efficient, V103I and I251L mutants fed with a hypercaloric diet increased their fasting glucose levels and adiposity similar to their wild-type littermates.

**Conclusions:** Altogether, our results demonstrate that mice carrying V103I and I251L MC4R mutations displayed gain-of-function phenotypes that were more evident in females. However, hypermorphic MC4R mutants were as susceptible as their control littermates to the obesogenic and diabetogenic effects elicited by a long-term hypercaloric diet, highlighting the importance of healthy feeding habits even under favorable genetic conditions.

**Highlights:**

- Identical single nucleotide substitutions to gain-of-function mutations of the human MC4R were introduced into the genome of inbred mice using CRISPR/Cas9 technology.
- Homozygous female carriers of V103I *Mc4r* alleles are shorter and display reduced adiposity, together with improved glucose homeostasis in both sexes.
- Homozygous females carrying the I251L mutation in the mouse MC4R also display shorter longitudinal length, decreased weight in all abdominal white fat pads and improved glucose clearance and insulin response.
- Gain-of-function mutations V103I and I251L of MC4R do not protect against the diabetogenic and adipogenic effects of a high-fat diet.

## 1. Introduction

Over the last three decades the entire world has witnessed a dramatic increase in the prevalence of overweight and obesity in children, teenagers and adults with an associated raise in chronic conditions such as type 2 diabetes (T2D), hypertension and cardiovascular disease [1,2]. Although mainly driven by the massive intake of ultra-processed edibles of poor nutritional value, the obesogenic contemporary human lifestyle does not affect all individuals equally, suggesting some level of genetic predisposition to develop or resist overweight and increased adiposity. Several genome-wide association studies performed during the last 15 years identified nearly 100 different loci associated with high body mass index (BMI), increased adiposity, high leptin levels or T2D [3–6]. However, most familial cases appear to be driven by several coexisting low-frequency polymorphic alleles, each of them contributing with minor effects and a small percentage to the overall phenotype. Notwithstanding, there are a few exceptions of human monogenic obesity syndromes including variants of genes involved in the central melanocortin system [7,8].

The melanocortin 4 receptor (MC4R) is a G protein-coupled receptor mainly expressed in the brain and autonomic sympathetic ganglia that is involved in the regulation of food intake and energy expenditure [9,10]. Loss-of-function mutations in the *MC4R* constitute the most common monogenic cases of early-onset extreme obesity in humans worldwide [11,12]. Individuals carrying non-synonymous MC4R variants display different levels of overweight while those carrying total loss of function mutations display voracious hyperphagia and early-onset extreme obesity, such as *Mc4r* null-allele mice that become extremely hyperphagic and obese since weaning [13].

Besides the existence of several hypomorphic alleles predisposing to obesity, some individuals carry non-synonymous MC4R variants that appear to protect against overweight and increased adiposity. The most frequent MC4R polymorphism reported to have a protective effect on obesity is V103I, with a carrier frequency of ∼2 to 4% [14,15]. *In vitro* studies performed in HEK293 cells stably transfected with a human MC4R clone carrying the V103I mutation showed a 2-fold decrease in the potency of the MC4R antagonist hAGRP(87–132) [16] suggesting diminished activity of endogenous AGRP in the blockade of MC4R necessary to promote food intake. Several case-control studies performed with individuals of European origin [17,18] and subsequent meta-analysis [14,19] showed that individuals carrying the V103I polymorphism have 2 to 18 % lower risk for obesity. Another large meta-analysis that included six East Asian studies and 31 studies of other ethnic groups, involving 19,822 obese cases and 35,373 non-obese controls showed that individuals with the V103I allele have 21% lower risk for obesity [20]. However, other studies performed with people of Chinese or Japanese descent showed no difference in obesity rates between*MC4R* 103V and 103I carriers [21,22]. Another non-synonymous polymorphism of MC4R, I251L, has also been reported to confer a 48% reduced risk for obesity in a meta-analysis on Europeans [19], in agreement with a previous study showing that this I251L substitution confers elevated MC4R-dependent cAMP levels in transfected cells [16]. In eight out of nine case-control studies performed in populations of European descent, the MC4R I251L polymorphism has been reproducibly associated with protection against obesity. A more recent study based on genetic and clinical data obtained from near half a million people of European-ancestry identified 61 non-synonymous MC4R variants including 4 gain of function alleles (V103I, I251L, I289L, I317V) characterized by enhanced β-arrestin recruitment rather that increased cAMP production when tested in transfected HEK293 cells [15]. Interestingly, carriers of these 4 gain-of-function variants showed significantly lower body mass index and lower risk of obesity, type 2 diabetes and coronary artery disease [15].

Although these studies suggest that gain of function variants of MC4R provide protection for increased adiposity and associated disorders, the multigenic nature underlying food intake and energy expenditure together with the great variability of the human genetic pool and people’s habits regarding food intake and physical activity also uncover the difficulty to rigorously test the functional association of a polymorphic allele with a particular behavior, physiological parameter or risk of disease. To directly challenge the hypothesis that some MC4R polymorphic variants protect against obesity we introduced the most common human gain of function mutations V103I and I251L into identical *Mc4r* coding positions of inbred laboratory mice which, together with a homogeneous environmental setting, provide an ideal experimental framework where to functionally test the effects of these variants in energy balance.

## 2. Materials and Methods

### 2.1. Multiple sequence alignment

Mouse and human MC4R amino acid sequences were aligned using ClustalW2 [23] using default parameters. UniProt accession numbers for the MCRs proteins are: mMC4R, P56450; mMC1R, Q01727; mMC2R, Q64326; mMC3R, P33033; mMC5R, P41149; hMC4R, P32245; hMC1R, Q01726; hMC2R, Q01718; hMC3R, P41968; hMC5R, P33032.

### 2.2. Imaging and patch clamp experiments in HEK293T cells

Mouse *Mc4r* sequences and the mutant versions V103I and I251L were amplified by PCR from mouse genomic DNA (see primers sequences in Table S1) and inserted into the *EcoRI* and *BamHI* sites of PRK6-YFP vector (IGF, Montpellier, France) and Sanger-sequenced (Macrogen). Imaging and patch clamp experiments were performed in HEK293T cells transfected with *wild-type Mc4r, Mc4r*^*V103I*^ or *Mc4r*^*I251L*^ tagged with YFP coding sequences (yellow fluorescent protein) and treated with Hoechst fluorescent dye to stain nuclei. 48 h after transfection, 1 mg/ml of CellMask™ Orange plasma membrane stain (ThermoFisher Scientific) was added to the culture medium for 1 min at 37°C and cells were then washed three times with PBS. Fluorescence photomicrographs were obtained with a Zeiss epifluorescence microscope using a pseudoconfocal Apotome module. Analyses of photomicrographs were performed with FIJI free software. *Wild-type* and mutant MC4Rs functionality was tested in HEK293T cells by means of MC4R agonist-evoked activation of inhibitory currents on type 2.2 voltage-gated calcium channels (CaV2.2) after bath application of the MC4/MC3 agonist melanotan II (MTII, Phoenix Pharmaceutical), as previously described [24]. Constitutive activity of the MCRs was assessed in HEK293T cells by means of their inhibitory effect on type 2.1 voltage-gated calcium channels (CaV2.1), as previously described [25].

### 2.3. Animal Care

Mice were housed in ventilated cages under controlled temperature and photoperiod (12-h light/12-h dark cycle, lights on from 7:00 AM to 7:00 PM), with tap water and laboratory chow containing 28.8% protein, 5.33% fat, and 65.87% carbohydrate available *ad libitum*. High caloric diet containing 23% protein, 23% fat and 56% carbohydrate was prepared with a mixture of laboratory chow and peanut butter fudge. All mouse procedures followed the Guide for the Care and Use of Laboratory Animals [26] and in agreement with the Institutional Animal Care and Use Committee of the Instituto de Investigaciones en Ingeniería Genética y Biología Molecular (INGEBI, CONICET).

### 2.4. Mutant mouse generation

Single point nonsynonymous coding mutations in *Mc4r* were introduced following a homology directed repair strategy based on the CRISPR/Cas9 system and a single stranded oligodeoxynucleotide (ssODN) microinjected in FVB/NJ mouse zygotes. Single guides were selected using crispr.mit.edu website of the Zhang Lab and CasOFFinder algorithm [27] according to proximity to the codons of interest and low off-targets (Table S2). Guides were generated by oligonucleotide hybridization (IDT, sequences available in Table S1) and sub- cloned in plasmid DR274 (Addgene, Plasmid 42250). sgRNAs were synthesized using MEGAshortscript T7 Transcription Kit (Ambion, AM1354). Cas9 mRNA was synthesized from plasmid MLM3613 (Addgene, Plasmid #42251) using mMESSAGEmMACHINE™ T7 Transcription Kit (Ambion, AM1344) and Poly(A) Tailing Kit (Ambion, AM1350). 93 nt ssODNs were purchased (Sigma, sequences available in Table S1). Cas9 mRNA (50 ng/µl for V103I mice and 80 ng/µl for I251L mice), sgRNA (50 ng/µl for V103I and 30 ng/µl for I251L) and ssODN (50 ng/µl for V103I and 30 ng/µl for I251L) were injected into the cytoplasm and pronucleus of one-cell FVB/NJ embryos (microinjection details in Table S3). Founder mice were analyzed by PCR using primers described in Table S1 and PCR products were subcloned in pGEM.T Easy Vector for sequencing (Promega). F1 and F2 mice were confirmed by PCR and by PCR products sequencing (primers available in Table S1). Strains carrying V130I or I251L MC4R mutations were selected and maintained in a FVB/NJ background.

### 2.5. *Mc4r* mRNA quantification by RT-qPCR

Whole mouse adult hypothalami were dissected, collected in ice-cold TriPure Isolation Reagent (Sigma, 11667165001) and stored at −80°C until RNA extraction, which was performed following the manufacturer’s instructions. RNA integrity was assessed by gel electrophoresis with clear 28S and 18s rRNA observed in an approximate 2:1 ratio. Quantification was performed using a Nanodrop and 260/280 and 260/230 nm ratios were checked to assess purity. 1 μg of RNA was treated with DNAse I (Ambion, AM2222) and used for first-strand cDNA synthesis, using High Capacity Reverse Transcription Kit with random primers (Applied Biosystems, 4368814). Primers were designed using the Primer 3 program (sequences are listed in Table S1). *Mc4r* mRNA was quantified using primers spanning the single *Mc4r* exon relative to internal control transcript *β-actin*. Samples were run in triplicate in an Applied Biosystems 7500 Real-Time PCR System machine using Power Up SYBR Green Master Mix (Applied Biosystems, A25472). Melt curves were analyzed to confirm specificity of the PCR product. Relative quantification was done by interpolating Ct values in standard curves or by 2-ΔΔCT.

### 2.6. Body weight, length and fat composition

Body weight was monitored weekly from weaning until 16-weeks of age. After that, mice were euthanized by cervical dislocation. After measuring the body longitudinal axial length from the nose to the base of the tail, mice were dissected and the inguinal, retroperitoneal and gonadal white fads removed and weighed, followed by the interscapular brown adipose tissue and the livers.

### 2.7. Food intake

13 week-old mice were individually housed with *ad libitum* access to chow food pellets. A week later, daily food intake was determined by subtracting the weight of food pellets placed on each cage at 6 PM by that obtained the following mornings at 10 AM.

### 2.8. Glucose and insulin tolerance tests

Fifteen week-old mice were fasted for 18 h (6 PM to 12 PM) previous to a glucose tolerance test or for 4 h (8 AM to 12 PM) for the insulin tolerance test. For basal fasting levels, blood samples were taken from the tail tip and glucose concentration measured with a One Touch R glucometer (LifeScan, Johnson & Johnson). Following a D-glucose (1 g/kg; Sigma, G5767) i.p. injection, blood samples were taken at 15, 30, 60, and 120 min. For the insulin tolerance test, insulin (1 IU/kg, Humulin R; Lilly; i.p.) was administered and blood samples taken at 15, 30, 60, and 120 min. The total area under the curve (AUC) was calculated using the trapezoidal rule.

### 2.9. Statistics

All data presented are the mean ± SEM and were analyzed using GraphPad Prism Software (version 5.01, 2007, GraphPad Software) or R Studio (version 1.1.456) by ANOVA. Post hoc pairwise comparisons between groups were performed by Holm-Sidak test unless stated. Normality of the distributions was assessed by Shapiro–Wilk test (p > 0.05), and the equality of the variance with the Bartlett’s or F test (p > 0.05). P values <0.05 were considered to be significant.

## 3. Results

### 3.1. Generation of mutant mice carrying V103I and I251L MC4R variants

The human *MC4R* is highly polymorphic in its coding region with up to 220 non-synonymous mutations identified within its 322 amino acids (Figure 1A). Despite the great number of human variants, *MC4R* is a highly conserved vertebrate gene with a remarkable identity at the amino acid sequence level among mammals. In particular, human and mouse MC4Rs are 94.3% identical (Figure 1B). The MC4R 103-Val residue is highly conserved not only between human and mice (Figure 1B) but also in all vertebrate Classes. The 251-Ile position, however, is an amniote (reptiles, birds and mammals) novelty that evolved from a leucine still present in extant fish and amphibians. Thus, the I251L gain of function mutation found in humans represents a regression to the vertebrate ancestral amino acid. Interestingly, all other human and mouse melanocortin receptors (MC1R, MC2R, MC3R and MC5R) also carry a highly conserved leucine in this position (Figure 1C) as all vertebrates do, with the only exception of teleost fish MC2R.

**Figure 1.**
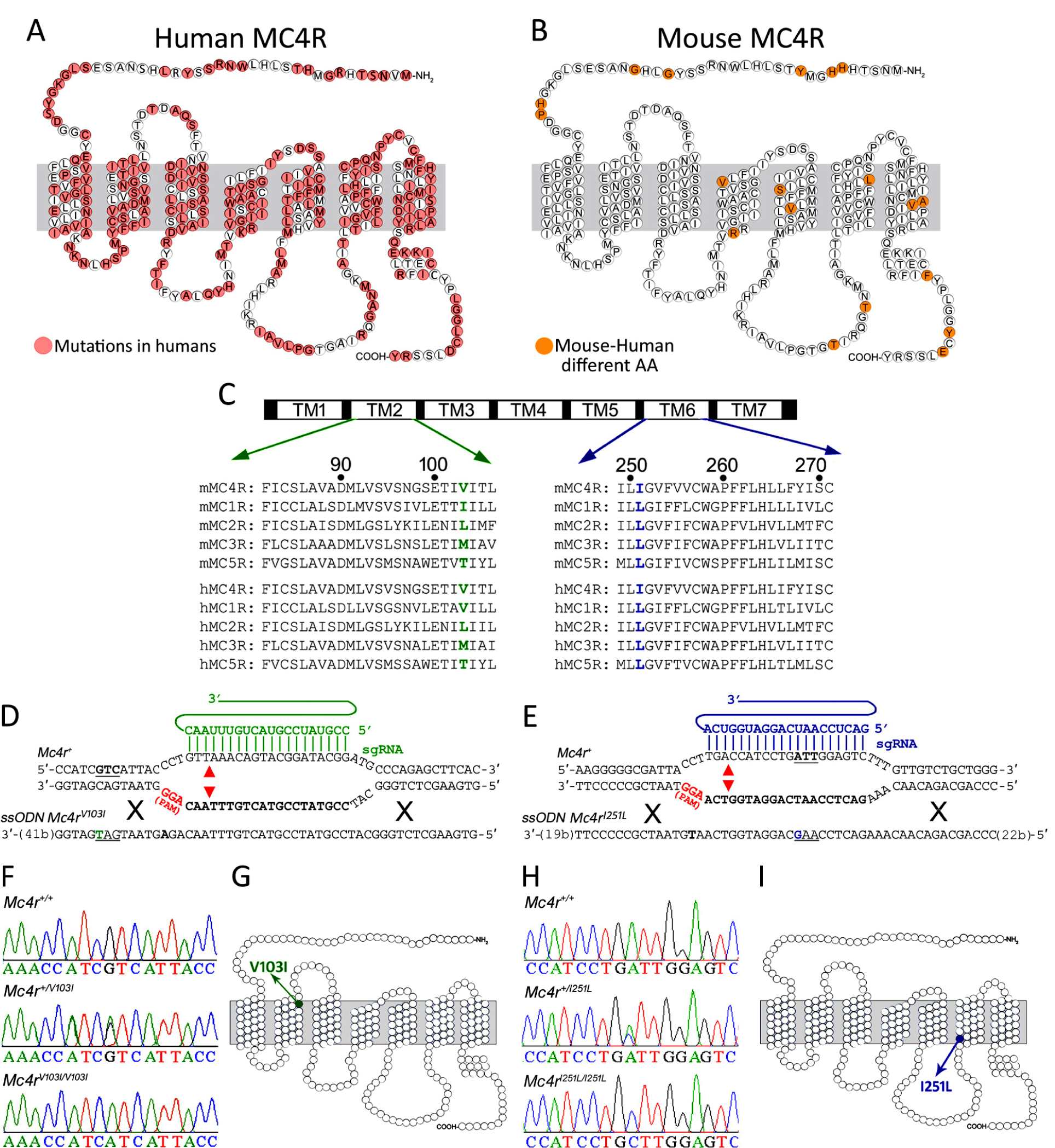
Generation of mutant mice carrying V103I and I251L MC4R variants. (A) Schematic of the human MC4R with all identified substitutions in red (mc4r.org.uk). (B) Schematic of the mouse MC4R. Non-conserved aminoacids between human and mouse are highlighted in orange. (C)Sequence alignment of the second and sixth transmembrane (TM) domains of mouse (top) and human (bottom) MC4R with MC1R, MC2R, MC3R and MC5R sequences. The 103-Val residue at TM2 is shown in green and the 251-Ile residue at TM6 is shown in blue. (D-E) Schematic of the targeting regions of the mouse *Mc4r* locus. sgRNAs driving Cas9-mediated cuts are shown in green for the V103I mutation and blue for the I251L mutation. Target sequences are shown in bold, the protospacer adjacent motifs (PAM) in red and the theoretical cut sites indicated with arrows. (D) As sODN donor containing a point mutation in codon 103 (green T) and a mutation in the PAM (bold A) was coinjected with a sgRNA and Cas9 mRNA into mouse zygotes to replace a coding Val to Ile (GTC to ATC) in the 103 residue. (E) Another ssODN donor was used to replace a coding Ile to Leu (ATT to CTT, shown in blue), in position 251. (F) Representative *Mc4r* PCR products confirming the c.307G>A point mutation in *Mc4r*^*V103I*^ alleles that predict a V103I mutation (G). (H) PCR products showing the c.751A>C point mutation in *Mc4r*^*I251L*^allelesthat predict a I251L substitution (I).

After assessing that mouse MC4R variants V103L and I251L are functional in transfected cells (Figure S1), we sought to investigate the *in vivo* functional effects of these two mutations. To this end, we generated two *knock-in* mutant mouse strains carrying the identical mutations found in the human population by means of CRISPR/Cas9-mediated gene-editing. For each mutation we microinjected FVB/NJ mouse zygotes with one sgRNA targeting *Mc4r* sequences close to the mutation together with a 93-nt ssODN donor carrying the identical nonsynonymous point mutations found in humans flanked by mouse *Mc4r* sequences (Tables S1 and S2) to promote homologous-directed repair (Figure 1D and 1E). DNA sequences at the target sites from all mutant F0 mice generated are shown in Table S4. *Mc4r* DNA sequences taken from F2 mice confirmed the c.307 G>A point mutation that converts the amino acid V103 into a coding isoleucine (Figure 1F and 1G), whereas a c.751 A>C point mutation changes amino acid I251 into a coding leucine (Figure 1H and 1I).

### 3.2. *Mc4r* ^***V103I/V103I***^ **mice exhibit gain of function phenotypes**

Heterozygous *Mc4r*^*+/V103I*^ breeding pairs displayed normal fertility rates and produced mice of the 3 genotypes at Mendelian ratios. *Mc4r*^*V103I/V103I*^ mice of both sexes and their *wild-type*(WT) littermates displayed identical body weight curves up to 16 weeks of age when fed *ad libitum* with normal chow (Figure 2A). At this age, daily food intake in *Mc4r*^*V103I/V103I*^ females and males was not different from their WT siblings (p=0.4 and 0.1, respectively; Figure 2B). In addition, mice of all genotypes and sexes displayed normal hypothalamic *Mc4r* mRNA levels (Figure 2C). However, 16 week-old *Mc4r*^*V103I/V103I*^ females showed shorter body length than their WT and heterozygous littermates (Figure 2D, left). This difference was not observed in the mutant males (Figure 2D, right). The calculated BMI (g/cm2) of 16 week-old mice showed no statistical differences for all genotypes and sexes. Despite their normal body weight, abdominal white adipose tissue (WAT) of 16 week-old *Mc4r*^*V103I/V103I*^ females weighed 40% less than that of WT littermates (Figure 2E, left), a difference that was not observed between WT and mutant males (Figure 2E, right). The lower abdominal WAT weight found in *Mc4r*^*V103I/V103I*^ females was better explained by a reduction in visceral WAT (Figure 2F) rather than inguinal WAT (Figure 2G), with a major effect found in the visceral gonadal fat pad (Figure 2H) and a non-significant reduction in retroperitoneal WAT (Figure 2I). The weight differences observed between the fat pads of *Mc4r*^*V103I/V103I*^ and WT littermates were female-specific and not found in males (Figure 2E-I). The weight of the interscapular brown adipose tissue (BAT) and the liver were normal in males and females of all genotypes (Figure 2J and 2K).

**Figure 2.**
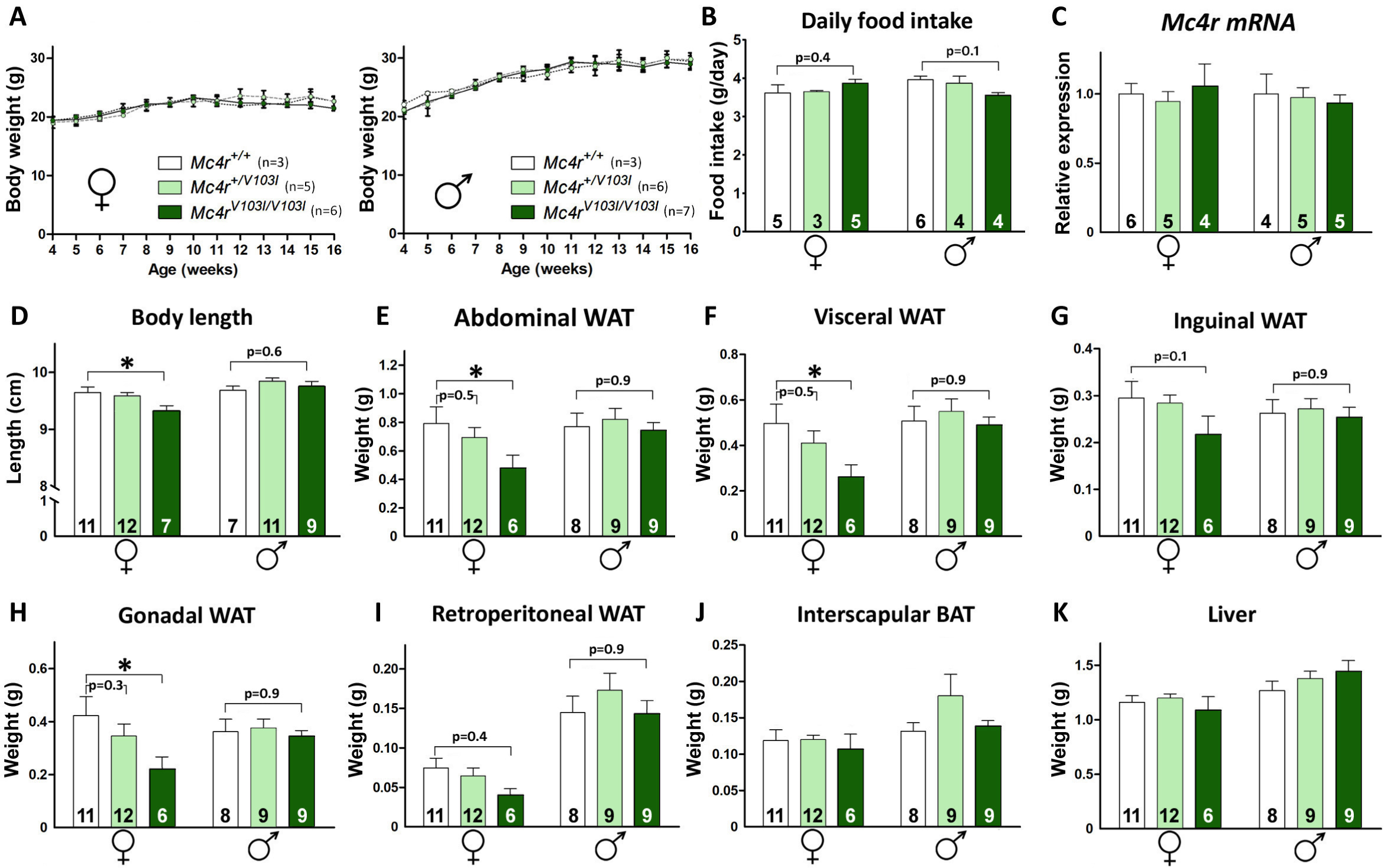
Gain of function phenotypes in *Mc4r*^***V103I/V103I***^**mice**. (A) Body weight curves of *Mc4r*^*+/V103I*^ and *Mc4r*^*V103I/V103I*^ females and males and their respective controls (n=3-7 per group). (B) Average daily food intake measured during 2 consecutive weeks at 14 weeks of age (n=3-6 per group). (C) Hypothalamic *Mc4r* mRNA levels in females and males of the 3 genotypes at 16 weeks of age (n=4-6 per group) assessed by qRT-PCR and expressed in arbitrary units. (D) Body length of 16 week-old females and males of the 3 genotypes (n=7-12 per group). (E-K) Determination of abdominal, visceral, inguinal, gonadal, and retroperitoneal white fat pad, interscapular brown fat pad and liver weights in 16-week-old female and male *Mc4r*^*+/V103I*^ and *Mc4r*^*V103I/V103I*^ mice and their respective controls (n=6-12 per group). Values represent the mean ± SEM. *P < 0.05 (Two-way ANOVA followed by Holm-Sidak test).

### 3.3. *Mc4r* ^***I251L/I251L***^ **mice also exhibit gain of function phenotypes**

Similar to *Mc4r*^*+/V103I*^ mice, heterozygous carriers of the I251L mutation were healthy and fertile. We raised a colony of *Mc4r*^*I251L/I251L*^ mice and WT littermates by mating heterozygote *Mc4r*^*+/I251L*^ mice and found that body weight curves up to 16 weeks of age in mutant mice of both sexes (Figure 3A). Daily food intake was indistinguishable between WT and homozygous mutant females (p=0.3) or males (p=0.6) (Figure 3B). In addition, hypothalamic levels of *Mc4r* mRNA were normal in all genotypes and sexes (Figure 3C). *Mc4r*^*I251L/I251L*^ females exhibited a shorter body length than their WT littermates (Figure 3D, left), a difference that was not found in males (Figure 3D, right). BMI of *Mc4r*^*I251L/I251L*^ mice is also normal in both sexes. The abdominal and visceral WAT were remarkably lighter in 16 week-old *Mc4r*^*I251L/I251L*^ females in comparison to what we observed in WT female littermates (Figure 3E and 3F). Different from the V103I heterozygote mutants, *Mc4r*^*+/I251L*^ females also showed reduced abdominal and visceral WAT weight compared to their WT female littermates (Figure 3E and 3F). Again, these differences were not found in males. The reduced fat accumulation was observed in the three white fat pads studied (inguinal, gonadal and retroperitoneal) of *Mc4r*^*I251L/I251L*^ females which displayed lower weights than those observed in their WT female littermates (Figure 3G, 3H and 3I). The visceral fat fads (gonadal and retroperitoneal) of heterozygous *Mc4r*^*+/I251L*^ females also showed to be lighter than in WT females (Figure 3H and 3I). As found with the V103I mutants, the weight of interscapular BAT and livers of *Mc4r*^*I251L/I251L*^ mice were normal in both sexes (Figure 3J and 3K).

**Figure 3.**
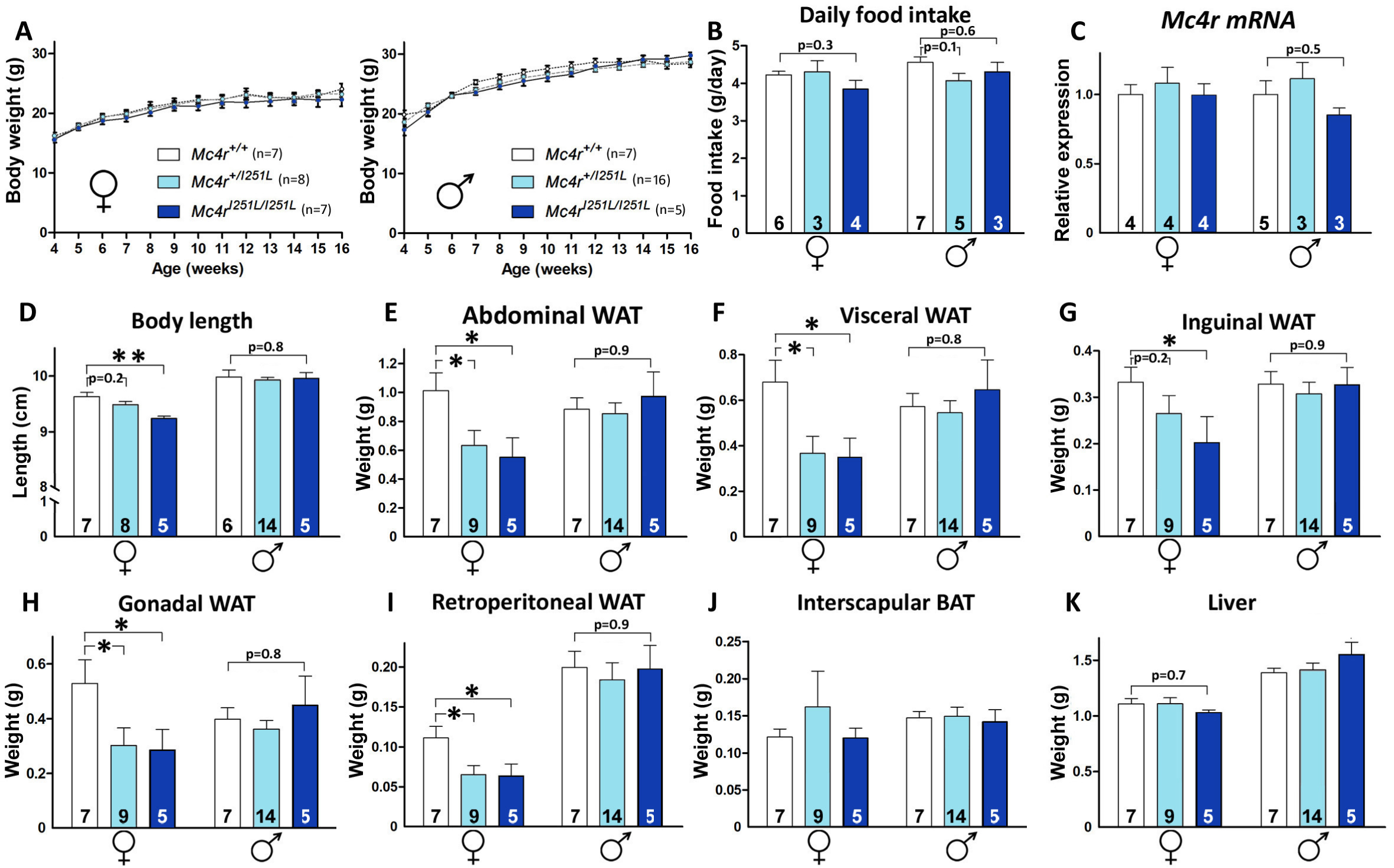
*Mc4r* ^***I251L/I251L***^ mice also exhibit gain of function phenotypes. (A) Body weight curves of *Mc4r*^*+/I251L*^ and *Mc4r*^*I251L/I251L*^ females and males and their respective controls (n=5-16 per group). (B) Average daily food intake measured during 2 consecutive weeks at 14 weeks of age (n=3-7 per group). (C) Hypothalamic *Mc4r* mRNA levels in *Mc4r*^*+/I251L*^ and *Mc4r*^*I251L/I251L*^ females and males at 16 weeks of age relative to controls (n=3-5 per group) assessed by qRT-PCR and expressed in arbitrary units. (D) Body length of 16 week-old females and males of the 3 genotypes (n=5-14 per group). (E-K) Determination of abdominal, visceral, inguinal, gonadal, and retroperitoneal white fat pad, interscapular brown fat pad and liver weights in 16-week-old female and male *Mc4r*^*+/I251L*^ and *Mc4r*^*I251L/I251L*^ mice and their respective controls (n=5-14 per group). Values represent the mean ± SEM. *P < 0.05, **P < 0.01 (Two-way ANOVA followed by Holm-Sidak test).

### 3.4. Mice carrying MC4R V103I and I251L mutations display more efficient glucose homeostasis

Since V103I and I251L human variants have been associated with lower odds of type 2 diabetes [15], we sought to determine whether the mutant mice carrying either of these two mutations have improved glucose homeostasis. *Mc4r*^*V103I/V103I*^ males and females showed normal fasting blood glucose levels (Figure 4A). Interestingly, female and male homozygous mutants showed improved glucose homeostasis evidenced by lower raises in blood glucose concentrations 30 and 60 min after receiving an intraperitoneal injection of D-glucose (1 g/kg) than the levels found in their *Mc4r*^*+/V103I*^ and WT littermates (Figure 4B and 4C). An insulin tolerance test performed in WT and V103I mice showed that insulin sensitivity is normal in all genotypes and sexes (Figure 4E and 4F).

**Figure 4.**
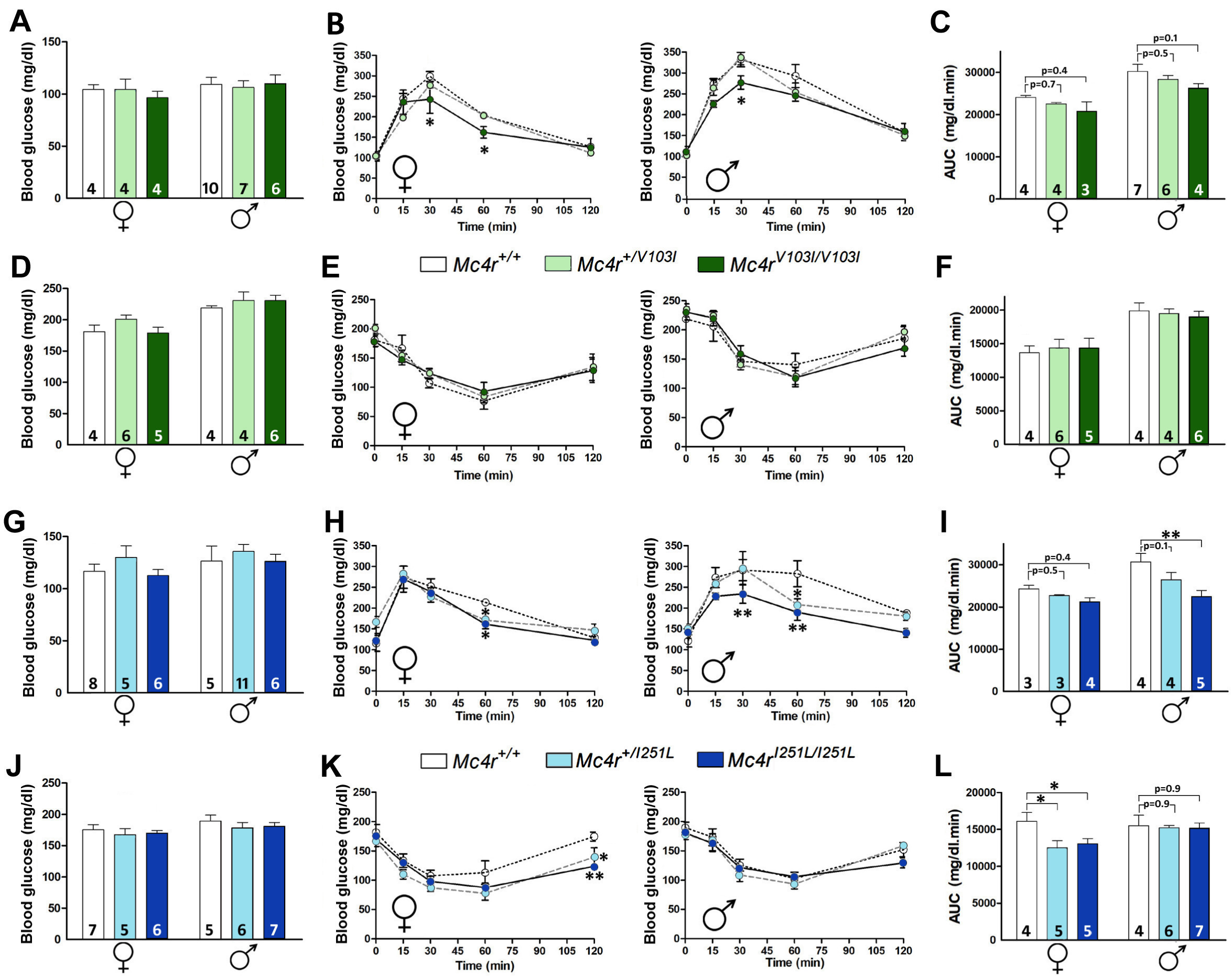
Mice carrying MC4R V103I and I251L mutations display more efficient glucose homeostasis. (A) Blood glucose concentration in 15-week-old WT, *Mc4r*^*+/V103I*^ and *Mc4r*^*V103I/V103I*^ female and male littermates after an 18 h fast (n=4-10 per group), (B) Glucose tolerance test (GTT) in mice receiving a 1 g/kg (i.p.) injection of D-glucose (n=3-7 per group; P: genotype effect of RMA *P< 0.05; Holm-Sidak) and, (C) the area under the curve (AUC). (D) Blood glucose concentration in 15-week-old WT,*Mc4r*^*+/V103I*^ and *Mc4r*^*V103I/V103I*^ female and male siblings after a 4 h fast (n=4-6 per group), (E) Insulin tolerance test (ITT) in mice receiving a 1 IU/kg (i.p.) injection of insulin (n=4-6 per group) and (F) AUC of E. (G) Blood glucose concentration in 15- week-old WT, *Mc4r*^*+/I251L*^ and *Mc4r*^*I251L/I251L*^ female and male littermates after an 18 h fast (n=5- 11 per group), (H) GTT (n=3-5 per group; P: genotype effect of RMA *p < 0.05; **p< 0.01; Holm- Sidak) and (I) AUC of H, **p <0.01; Two-way ANOVA, Holm-Sidak. (J) Blood glucose concentration in 15-week-old WT,*Mc4r*^*+/I251L*^ and *Mc4r*^*I251L/I251L*^ female and male siblings after a 4 h fast (n=5-7 per group), (K) ITT (n=4-7 per group; p: genotype effect of RMA *p < 0.05; **p < 0.01; Holm-Sidak) and (L) AUC of K, *P<0.05; Two-way ANOVA, Holm-Sidak. Values represent the mean ± SEM.

Similar to V103I mutants, *Mc4r*^*I251L/I251L*^ females and males also showed normal fasting glycaemia and improved glucose clearing when challenged in a glucose tolerance test (GTT; Figure 4H and 4I). Interestingly, *Mc4r*^*I251L/I251L*^ females, but not males, showed improved insulin response when challenged in an insulin tolerance test (Figure 4K and 4L). Altogether, these results show that MC4R V103I and I251L mutations lead to improved glucose homeostasis suggesting a more efficient insulin response.

### 3.5. V1O3I and I251L MC4R mutations do not protect against a high-fat diet

Given that humans carrying gain of function *MC4R* variants were associated with lower BMI, obesity and T2D [15], and that *Mc4r*^*V103I/V103I*^ and *Mc4r*^*I251L/I251L*^ female mice showed reduced WAT adiposity (Figure 2 and 3), we decided to determine whether the V103I and I251L alleles would protect from the hyperglycemic and obesogenic effects normally elicited in FVB/N mice after a long-term high fat diet (HFD) [28,29]. After weaning, mice were fed *ad libitum* with a normal diet (ND) for two weeks and then switched to a HFD for 11 more weeks. At 16 weeks of age, *Mc4r*^*V103I/V103I*^ females and their WT littermates fed on a HFD were heavier than those receiving a ND (Figure 5A, left). However, no differences were found among genotypes receiving either ND or HFD (Figure 5A). Similar results were found in males (Figure 5A, right). The increased body weight is largely due to an exaggerated fat deposition in subcutaneous (inguinal; Figure 5B) and visceral (retroperitoneal and gonadal) white fat pads (Figure 5C and 5D) as well as in interscapular BAT (Figure 5E). The increased adiposity elicited by a long-term HFD was not different in any mouse genotypes or gender. In contrast to what we observed in all fat pads analyzed, the weights of the livers of mice receiving HFD were not statistically different from those measured in mice fed with a ND across all genotypes and genders. We also found that mice fed on a HFD exhibited a remarkable increase in fasting blood glucose levels that was similar across all genotypes and in both sexes (Figure 5F). In agreement with increased adiposity and higher fasting glucose levels, mice receiving a HFD showed impaired glucose clearing when challenged in a GTT. This HFD-induced deficit was similarly found in mice of all genotypes and sexes (Figure 5G and 5H). Although being hyperglycemic, mice of all genotypes and sexes fed on a HFD displayed normal responses to an insulin challenge that were no different from those observed in mice receiving a ND (Figure 5I and 5J).

**Figure 5.**
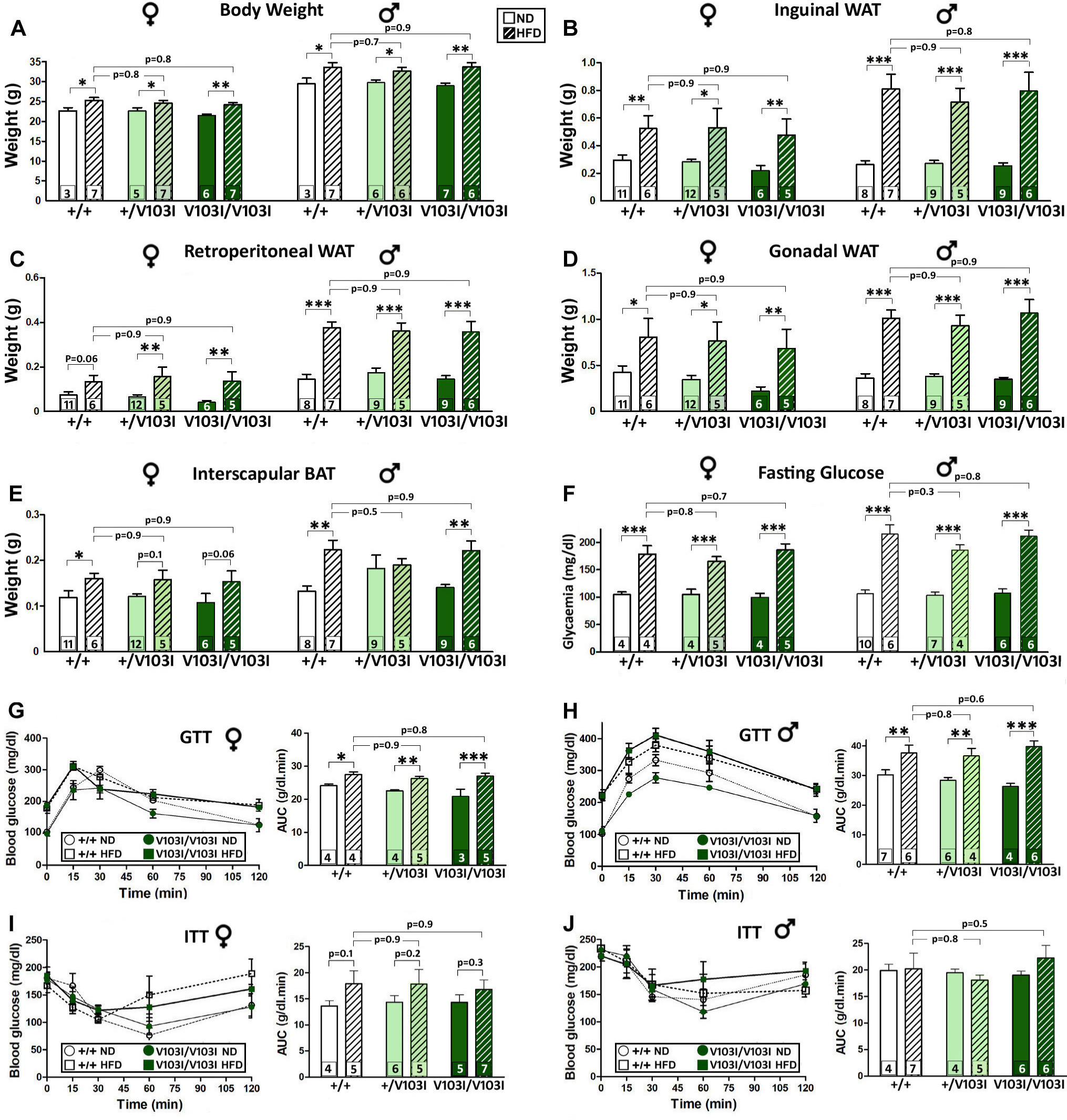
The V1O3I MC4R mutation does not protect against a high-fat diet. Metabolic phenotype of *Mc4r*^*+/V103I*^ and *Mc4r*^*V103I/V103I*^ females and males and their respective WT controls under the effect of a 3 month-high fat diet (HFD) compared to a normal diet (ND). (A) Body weight of 16 week-old mice (n=3-7 per group). (B-E) Determination of inguinal (B), retroperitoneal (C) and gonadal (D) white fat pads and interscapular brown fat pad (E) of 16 week-old mice (n=5-12 per group). (F) Blood glucose concentration after 18 h fasting in 15 week-old mice (n=4-10 per group). (G-H) GTT using 1 g/kg D-glucose i.p. injection in 15 week- old mice fasted for 18 h. Bars represent the area under the curve (n=3-7 per group). (I-J) ITT using 1 IU/kg human recombinant insulin (i.p.) in 15 week-old mice of both sexes fasted for 4 h. Bars represent the area under the curve (n=5-7 per group). *P< 0.05, **P < 0.01, ***P< 0.001; Two-way ANOVA, Holm-Sidak. Values represent the mean ± SEM.

Analysis of mice carrying the I251L mutation in MC4R and fed on a HFD yielded very similar results to those described above for carriers of V103I alleles, including higher total body weight at 16 weeks of age (Figure 6A), increased weight in white fat pads (Figure 6B and 6D), elevated fasting blood glucose levels (Figure 6F), and impaired glucose clearing when challenged in a GTT (Figure 6G and 6H). Thus, in all these parameters we did not observe any difference between WT and *Mc4r*^*I251L/I251L*^ females or males fed with a HFD. We have also detected a few differences in the I251L group relative to the V103I cohort that included an impaired response to insulin in females carrying one or two I251L alleles fed with a HFD relative to those receiving a ND (Figure 6I) and in males of all genotypes (Figure 6J). Altogether, these results indicate that although homozygous carriers of V103I or I251L mutations in the MC4R exhibit reduced adiposity and improved glucose metabolism they are not protected against the obesogenic and diabetogenic effects elicited by a HFD.

**Figure 6.**
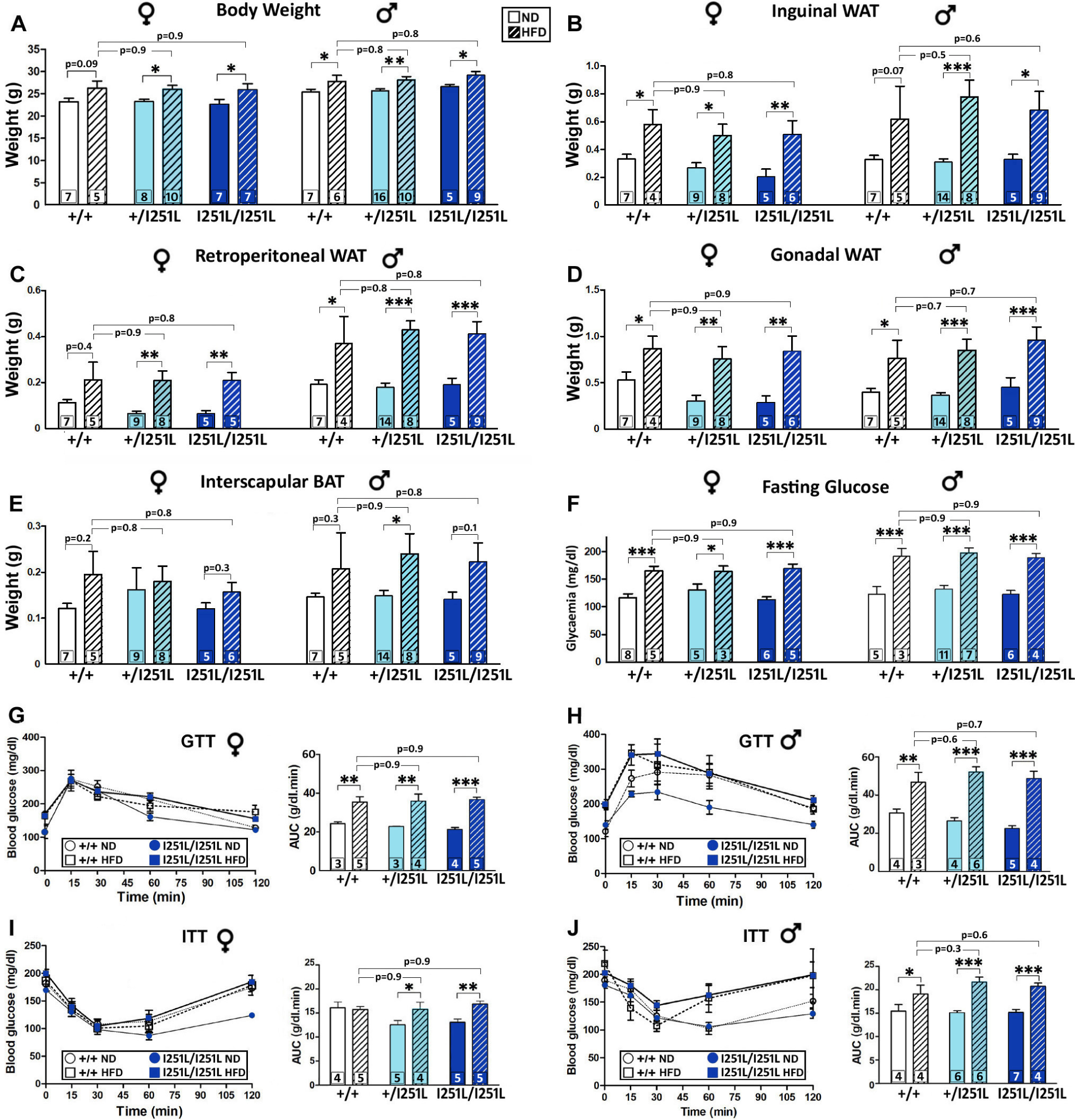
The I251L MC4R mutation does not protect against a high-fat diet. Metabolic phenotype of *Mc4r*^*+/I251L*^ and *Mc4r*^*I251L/I251L*^ females and males and their respective WT control littermates under the effect of a 3 month-high fat diet (HFD) compared to a normal diet (ND). (A) Body weight of 16 week-old mice (n=5-16 per group). (B-E) Determination of inguinal (B), retroperitoneal (C) and gonadal (D) white fat pads and interscapular brown fat pad (E) of 16 week-old mice (n=4-14 per group). (F) Blood glucose concentration after 18 h fasting in 15 week-old mice (n=3-11 per group). (G-H) GTT using 1 g/kg D-glucose i.p. injection in 15 week- old females and males fasted 18 h. Bars represent the area under the curve (n=3-6 per group). (I-J) ITT using 1 IU/kg human recombinant insulin (i.p.) in 15 week-old females and males fasted for 4 h. Bars represent the area under the curve (n=4-7 per group). *P < 0.05, **P < 0.01, ***P< 0.001; Two-way ANOVA, Holm-Sidak. Values represent the mean ± SEM.

## 4. Discussion

The search for the genetic bases underlying a higher predisposition for obesity and T2D has been boosted in the last two decades by several genome-wide studies involving a large number of case-control groups in different countries [3,4]. Although the statistical power of these studies allowed the association of more than 100 loci with obesity-related phenotypes, each identified gene showed a relatively small effect on disease risk. Given the incalculable loss of years and quality of life of the patients and the enormous economic consequences of the obesity pandemic, the low predictive value of the vast majority of these *loci* for assessing an individual’s risk is disappointing. One of the few exceptions, however, is *MC4R* since its nonsynonymous coding mutations are responsible for severe early-onset obesity [8,30] and SNPs in its 3’ flanking region have been associated with elevated BMI [31]. Recent population studies have also found nonsynonymous coding mutations in MC4R that appear to protect from obesity and T2D [15]. Although these human mutations displayed hypermorphic (gain of function) outcomes when tested *in vitro* they have never been experimentally tested *in vivo*. Because the genetic programs, neuroanatomical circuits and physiological regulation of the human and mouse central melanocortin systems are highly similar, mouse models designed to study the effects of orthologous human mutations have high construct and predictive value providing a useful platform where to analyze the functional effects of such mutations in a live mammal.

Thus, in this study we challenged the hypothesis that the V103I and I251L mutations in the mouse MC4R protects from increased adiposity and T2D, as has been suggested in association studies of human carriers of either one of these orthologous gain-of-function mutations [14,15,19]. To this end, we generated and studied mutant mice carrying the same nucleotide substitutions yielding V103I and I251L MCR4 alleles. In particular we show that: 1) Homozygous females carrying the V103I mutation display shorter longitudinal length and decreased abdominal white fat mainly due to a large reduction in gonadal fat; 2) Homozygous females carrying the I251L mutation also display shorter longitudinal length and decreased weight of all abdominal white fat pads examined: gonadal, retroperitoneal and inguinal; 3) Homozygous *Mc4r*^*V103I/V103I*^ and *Mc4r*^*I251L/I251L*^ mice of both sexes show improved glucose homeostasis when challenged with a GTT; 4) *Mc4r*^*I251L/I251L*^ females, but not males, showed improved insulin response when challenged in an insulin tolerance test; 5) the MC4R V103I and I251L mutations do not protect against the diabetogenic and adipogenic effects of a HFD as evidenced by a similar increase in fasting glucose levels and augmented adiposity in *Mc4r*^*V103I/V103I*^ and *Mc4r*^*I251L/I251L*^ mice of both sexes in comparison to their WT littermates. Altogether, the phenotypes observed in mice carrying *Mc4r* alleles with homozygous V103I or I251L substitutions are compatible with gain-of-function mutations, in contrast with loss-of-function MC4R mutants that display increased adiposity, longer stature and T2D [11-13]. Further studies will be needed to determine potential differences in respiratory exchange ratio, energy expenditure and distribution of fat deposition in the mutant mice.

Homozygous mutant females carrying V103I or I251L *Mc4r* alleles display shorter longitudinal growth, a phenotype also found in a recent association study performed in children carrying V103I MC4R variants [32]. Animal longitudinal length and human height are highly heritable polygenic traits, and although many genes contribute with a small percentage to the overall variation of human height there are some monogenic examples that produce relatively large effects, including MC4R. In fact, MC4R loss-of-function mutations in the human population have been linked to increased stature [33,34] and a similar phenotype was described in MC4R null-allele mutant mice [13]. Given the importance of insulin in body growth [35] and that MC4R stimulation diminishes insulin release in mice [36], it is tempting to speculate that hypermorphic V103I and I251L MC4Rs lead to a reduction in insulin levels and, consequently, to shorter longitudinal body length.

A noticeable outcome of our study is that the reduced adiposity, shorter axial length and improved glucose metabolism observed in V103I and I251L MC4R mutants were much more pronounced in females than in males. Body fat distribution and energy balance are sexually dimorphic, probably because most metabolic traits are highly influenced by sexual hormones. For example, male mice display higher food intake and energy expenditure even when normalized by their larger body mass and also gain more body weight than females when fed on a hypercaloric diet. Females, in turn, have more POMC neurons in the arcuate nucleus of the hypothalamus [37] and are more sensitive to the anorexigenic effects of central leptin [38,39]. In agreement, sexually dimorphic phenotypes are also commonly found in animal models used to study the effects of gene mutations in the control of energy balance and metabolism. For example, we previously showed that extremely obese mice lacking hypothalamic *Pomc* considerably improved their body weight and adiposity only in females once *Pomc* expression is reestablished, whereas males were more resistant to regain their normal weight [40]. In addition, *Pomc* rescue restricted to 5-HT2C positive arcuate neurons reestablished normal levels of physical activity, energy expenditure and body weight only in male mice [41]. Regarding *MC4R*, female mice lacking this receptor exhibit an even greater increase in body weight than males [13]. In humans, also, SNP rs17782313 has been linked to obesity and increased T2D risk only in women [42–44] whereas several loss of function human variants showed a greater increase in BMI in females than in males [45]. Another study showed that women carrying the gain of function mutation V103I displayed a more pronounced reduction in BMI than men [46]. Altogether, these studies suggest that females are more responsive to loss or gain of function mutations of this receptor.

The MC4R exhibits a great level of genetic variation in the human population (Figure 1A). For example, a recent association study performed in 452,300 individuals of European ancestry taken from the UK Biobank detected 61 non-synonymous polymorphic variants including 46 missense mutations that yielded loss and/or gain of function phenotypes when tested in transfected HEK293 cells for cAMP production or the recruitment of β-arrestin-2 [15]. While loss-of-function variants identified in this study were associated with higher odds of obesity, T2D and coronary artery disease; individuals carrying gain of function variants showed lower odds to develop these conditions. In particular, the gain of function variants V103I and I251L are present in the UK Biobank population at relatively high frequencies (2.0% and 1.3%, respectively), in quite contrast to all other variants that showed an allele frequency lower than 0.07%. It is tempting to speculate that these two alleles are maintained at relatively higher frequencies because they provide some level of protection against environmental conditions of excessive overconsumption of hypercaloric edibles.

Interestingly, the melanocortin 1 receptor gene (*MC1R*), paralog and ancestor of *MC4R*, has also shown to be polymorphic in its coding region and to produce heritable gain of function variation with functional adaptive consequences [47,48]; as found in several wild mouse populations that underwent an evolutionary darkening of their dorsal pelage after sandy rock terrains transitioned to black cooled lava flow lands following activity of nearby volcanoes [49].The newly established darker landscape was likely to exert selective pressure for the fixation of novel *Mc1r* mutations that promoted the production of dark eumelanin-stained hairs restoring camouflage adaptation against visual predators such as hunting owls. Thus, *MC4R* and *MC1R* are likely to share high levels of evolvability, defined as an ability to produce heritable and adaptive phenotypic variation driven by environmental changes [50]. While the thrifty gene theory suggests that genetic programs controlling energy balance evolved to adapt to extensive periods of food scarcity [51], such conditions are no longer present in vast sectors of modern human societies allowing hypermorphic variants of MC4R to coexist in relatively high frequencies within the current genetic pool and even provide some protection against the deleterious consequences of current obesogenic diets. Although the V103I and I251L mutant mice studied here showed to be leaner and metabolically more efficient when eating normal chow, they failed to overcome the negative consequences of a hypercaloric diet, highlighting the importance of healthy feeding habits even under favorable genetic conditions.

## Acknowledgments

The authors thank Marta Treimun and Carolina Álvarez for excellent technical work. This work was supported by Agencia Nacional de Promoción Científica y Tecnológica, PICT2016-1742. (M.R.). D.R. received a doctoral fellowship from CONICET.

## Conflict of interest

The authors declare no conflict of interest.

## Authors contributions

D.R. and M.R. designed this study; D.R., C.M. and J.R. conducted the research; J.R. and M.R. contributed new reagents and analytic tools; D.R., C.M., J.R. and M.R. analyzed the data and prepared the figures; M.R. wrote the paper; and D.R., C.M., J.R. and M.R. revised the paper.

## Appendix A. Supplementary Data

Supplemental Data include one figure.

## Appendix B. Supplementary Materials

Supplemental Materials include four tables.

## Notes

### Competing Interest Statement

The authors have declared no competing interest.

## References

[1] 2015. Preventing and Managing the Global Epidemic of Obesity. World Health Organization.

[2] James, W.P.T., 2018. Obesity: A global public health challenge. Clinical Chemistry 64(1): 24–9, Doi: 10.1373/clinchem.2017.273052.

[3] Thorleifsson, G., Walters, G.B., Gudbjartsson, D.F., Steinthorsdottir, V., Sulem, P., Helgadottir, A., et al., 2009. Genome-wide association yields new sequence variants at seven loci that associate with measures of obesity. Nature Genetics 41(1): 18–24, Doi: 10.1038/ng.274.

[4] Wheeler, E., Huang, N., Bochukova, E.G., Keogh, J.M., Lindsay, S., Garg, S., et al., 2013. Genome-wide SNP and CNV analysis identifies common and low-frequency variants associated with severe early-onset obesity. Nature Genetics 45(5): 513–7, Doi: 10.1038/ng.2607.

[5] Gaulton, K.J., Ferreira, T., Lee, Y., Raimondo, A., Mägi, R., Reschen, M.E., et al., 2015. Genetic fine mapping and genomic annotation defines causal mechanisms at type 2 diabetes susceptibility loci. Nature Genetics 47(12): 1415–25, Doi: 10.1038/ng.3437.

[6] Loos, R.J.F., 2012. Genetic determinants of common obesity and their value in prediction. Best Practice and Research: Clinical Endocrinology and Metabolism: 211–26, Doi: 10.1016/j.beem.2011.11.003.

[7] Krude, H., Biebermann, H., Luck, W., Horn, R., Brabant, G., Grüters, A., 1998. Severe early-onset obesity, adrenal insufficiency and red hair pigmentation caused by POMC mutations in humans. Nature Genetics 19(2): 155–7, Doi: 10.1038/509.

[8] Vaisse, C., Clement, K., Guy-Grand, B., Froguel, P., 1998. A frameshift mutation in human MC4R is associated with a dominant form of obesity [2]. Nature Genetics 20(2): 113–4, Doi: 10.1038/2407.

[9] Balthasar, N., Dalgaard, L.T., Lee, C.E., Yu, J., Funahashi, H., Williams, T., et al., 2005. Divergence of Melanocortin Pathways in the Control of Food Intake and Energy Expenditure. Cell 123(3): 493–505, Doi: 10.1016/j.cell.2005.08.035.

[10] Krashes, M.J., Lowell, B.B., Garfield, A.S., 2016. Melanocortin-4 receptor-regulated energy homeostasis. Nature Neuroscience 19(2): 206–19, Doi: 10.1038/nn.4202.

[11] Vaisse, C., Clement, K., Durand, E., Hercberg, S., Guy-Grand, B., Froguel, P., 2000. Melanocortin-4 receptor mutations are a frequent and heterogeneous cause of morbid obesity. Journal of Clinical Investigation 106(2): 253–62, Doi: 10.1172/JCI9238.

[12] Farooqi, I.S., Keogh, J.M., Yeo, G.S.H., Lank, E.J., Cheetham, T., O’Rahilly, S., 2003. Clinical spectrum of obesity and mutations in the melanocortin 4 receptor gene. The New England Journal of Medicine 348(12): 1085–95, Doi: 10.1056/NEJMoa022050.

[13] Huszar, D., Lynch, C.A., Fairchild-Huntress, V., Dunmore, J.H., Fang, Q., Berkemeier, L.R., et al., 1997. Targeted disruption of the melanocortin-4 receptor results in obesity in mice. Cell, Doi: 10.1016/S0092-8674(00)81865-6.

[14] Young, E.H., Wareham, N.J., Farooqi, S., Hinney, A., Hebebrand, J., Scherag, A., et al., 2007. The V103I polymorphism of the MC4R gene and obesity: Population based studies and meta-analysis of 29 563 individuals. International Journal of Obesity, Doi: 10.1038/sj.ijo.0803609.

[15] Lotta, L.A., Mokrosiński, J., Mendes de Oliveira, E., Li, C., Sharp, S.J., Luan, J., et al., 2019. Human Gain-of-Function MC4R Variants Show Signaling Bias and Protect against Obesity. Cell 177(3): 597–607.e9, Doi: 10.1016/j.cell.2019.03.044.

[16] Xiang, Z., Litherland, S. a., Sorensen, N.B., Proneth, B., Wood, M.S., Shaw, A.M., et al., 2006. Pharmacological characterization of 40 human melanocortin-4 receptor polymorphisms with the endogenous proopiomelanocortin-derived agonists and the agouti-related protein (AGRP) antagonist. Biochemistry 45(23): 7277–88, Doi: 10.1021/bi0600300.

[17] Gotoda, T., Scott, J., Aitman, T.J., 1997. Molecular screening of the human melanocortin-4 receptor genelll: identification of a missense variant showing no association with obesity, plasma glucose, or insulin 4: 976–9.

[18] Geller, F., Reichwald, K., Dempfle, A., Illig, T., Vollmert, C., Herpertz, S., et al., 2004. Melanocortin-4 receptor gene variant I103 is negatively associated with obesity. American Journal of Human Genetics 74(3): 572–81, Doi: 10.1086/382490.

[19] Stutzmann, F., Vatin, V., Cauchi, S., Morandi, A., Jouret, B., Landt, O., et al., 2007. Non-synonymous polymorphisms in melanocortin-4 receptor protect against obesity: The two facets of a Janus obesity gene. Human Molecular Genetics, Doi: 10.1093/hmg/ddm132.

[20] Wang, D., Ma, J., Zhang, S., Hinney, A., Hebebrand, J., Wang, Y., et al., 2010. Association of the MC4R V103I polymorphism with obesity: A chinese case-control study and meta-analysis in 55,195 individuals. Obesity, Doi: 10.1038/oby.2009.268.

[21] Ohshiro, Y., Sanke, T., Ueda, K., Shimajiri, Y., Nakagawa, T., Tsunoda, K., et al., 1999. Molecular scanning for mutations in the melanocortin-4 receptor gene in obese/diabetic Japanese. Annals of Human Genetics 63(6): 483–7, Doi: 10.1046/j.1469-1809.1999.6360483.x.

[22] Wang, C.L., Liang, L., Wang, H.J., Fu, J.F., Hebebrand, J., Hinney, A., 2006. Several mutations in the melanocortin 4 receptor gene are associated with obesity in Chinese children and adolescents. Journal of Endocrinological Investigation 29(10): 894–8, Doi: 10.1007/BF03349193.

[23] Larkin, M.A., Blackshields, G., Brown, N.P., Chenna, R., Mcgettigan, P.A., McWilliam, H., et al., 2007. Clustal W and Clustal X version 2.0. Bioinformatics 23(21): 2947–8, Doi: 10.1093/bioinformatics/btm404.

[24] Agosti, F., López Soto, E.J., Cabral, A., Castrogiovanni, D., Schioth, H.B., Perelló, M., et al., 2014. Melanocortin 4 receptor activation inhibits presynaptic N-type calcium channels in amygdaloid complex neurons. European Journal of Neuroscience 40(5): 2755–65, Doi: 10.1111/ejn.12650.

[25] Agosti, F., Cordisco Gonzalez, S., Martinez Damonte, V., Tolosa, M.J., Di Siervi, N., Schioth, H.B., et al., 2017. Melanocortin 4 receptor constitutive activity inhibits L-type voltage-gated calcium channels in neurons. Neuroscience 346(January): 102–12, Doi: 10.1016/j.neuroscience.2017.01.007.

[26] Nih., Od., Oer., Olaw., 2011. GUIDE LABORATORY ANIMALS FOR THE CARE AND USE OF Eighth Edition Committee for the Update of the Guide for the Care and Use of Laboratory Animals Institute for Laboratory Animal Research Division on Earth and Life Studies.

[27] Bae, S., Park, J., Kim, J.S., 2014. Cas-OFFinder: A fast and versatile algorithm that searches for potential off-target sites of Cas9 RNA-guided endonucleases. Bioinformatics 30(10): 1473–5, Doi: 10.1093/bioinformatics/btu048.

[28] Montgomery, M.K., Hallahan, N.L., Brown, S.H., Liu, M., Mitchell, T.W., Cooney, G.J., et al., 2013. Mouse strain-dependent variation in obesity and glucose homeostasis in response to high-fat feeding. Diabetologia 56(5): 1129–39, Doi: 10.1007/s00125-013-2846-8.

[29] Nascimento-Sales, M., Fredo-da-Costa, I., Borges Mendes, A.C.B., Melo, S., Ravache, T.T., Gomez, T.G.B., et al., 2017. Is the FVB/N mouse strain truly resistant to diet-induced obesity? hysiological Reports 5(9), Doi: 10.14814/phy2.13271.

[30] Yeo, G.S.H., Farooqi, I.S., Aminian, S., Halsall, D.J., Stanhope, R.G., O’Rahilly, S., 1998. A frameshift mutation in MC4R associated with dominantly inherited human obesity. Nature Genetics 20(2): 111–2, Doi: 10.1038/2404.

[31] Loos, R.J.F., Loos, R.J.F., Lindgren, C.M., Lindgren, C.M., Li, S., Li, S., et al., 2008. Common variants near MC4R are associated with fat mass, weight and risk of obesity. Nature Genetics 40(6): 768–75, Doi: 10.1038/ng.140.Common.

[32] Herrfurth, N., Volckmar, A.L., Peters, T., Kleinau, G., Müller, A., Cetindag, C., et al., 2018. Relevance of polymorphisms in MC4R and BDNF in short normal stature. BMC Pediatrics 18(1): 5–9, Doi: 10.1186/s12887-018-1245-1.

[33] Martinelli, C.E., Keogh, J.M., Greenfield, J.R., Henning, E., Van Der Klaauw, A.A., Blackwood, A., et al., 2011. Obesity due to melanocortin 4 receptor (MC4R) deficiency is associated with increased linear growth and final height, fasting hyperinsulinemia, and incompletely suppressed growth hormone secretion. Journal of Clinical Endocrinology and Metabolism 96(1): 181–8, Doi: 10.1210/jc.2010-1369.

[34] Vollbach, H., Brandt, S., Lahr, G., Denzer, C., Von Schnurbein, J., Debatin, K.M., et al., 2017. Prevalence and phenotypic characterization of MC4R variants in a large pediatric cohort. International Journal of Obesity 41(1): 13–22, Doi: 10.1038/ijo.2016.161.

[35] Hill, D.J., Milner, R.D.G., 1985. Insulin as a Growth Factor. vol. 19.

[36] Fan, W., Dinulescu, D.M., Butler, A.A., Zhou, J., Marks, D.L., Cone, R.D., 2000. The central melanocortin system can directly regulate serum insulin levels. Endocrinology 141(9): 3072–9, Doi: 10.1210/endo.141.9.7665.

[37] Wang, C., He, Y., Xu, P., Yang, Y., Saito, K., Xia, Y., et al., 2018. TAp63 contributes to sexual dimorphism in POMC neuron functions and energy homeostasis. Nature Communications 9(1): 1–11, Doi: 10.1038/s41467-018-03796-7.

[38] Clegg, D.J., Riedy, C.A., Smith, K.A.B., Benoit, S.C., Woods, S.C., 2003. Differential sensitivity to central leptin and insulin in male and female rats. Diabetes 52(3): 682–7, Doi: 10.2337/diabetes.52.3.682.

[39] Clegg, D.J., Brown, L.M., Woods, S.C., Benoit, S.C., 2007. Erratum: Gonadal hormones determine sensitivity to central leptin and insulin (Diabetes (2006) 55 (978-987)). Diabetes 56(10): 2649, Doi: 10.2337/db07-er10.

[40] Bumaschny, V.F., Yamashita, M., Casas-Cordero, R., Otero-Corchón, V., de Souza, F.S.J., Rubinstein, M., et al., 2012. Obesity-programmed mice are rescued by early genetic intervention. Journal of Clinical Investigation 122(11): 4203–12, Doi: 10.1172/JCI62543.

[41] Burke, L.K., Doslikova, B., D’Agostino, G., Greenwald-Yarnell, M., Georgescu, T., Chianese, R., et al., 2016. Sex difference in physical activity, energy expenditure and obesity driven by a subpopulation of hypothalamic POMC neurons. Molecular Metabolism 5(3): 245–52, Doi: 10.1016/j.molmet.2016.01.005.

[42] Horstmann, A., Kovacs, P., Kabisch, S., Boettcher, Y., Schloegl, H., Tönjes, A., et al., 2013. Common Genetic Variation near MC4R Has a Sex-Specific Impact on Human Brain Structure and Eating Behavior. PLoS ONE 8(9): 14–6, Doi: 10.1371/journal.pone.0074362.

[43] Liu, G., Zhu, H., Lagou, V., Gutin, B., Barbeau, P., Treiber, F.A., et al., 2014. Common variants near MC4R are associated with general and visceral adiposity in European- and African-American youth. J Pediatr 156(4): 598–605, Doi: 10.1016/j.jpeds.2009.10.037.Common.

[44] Renström, F., Payne, F., Nordström, A., Brito, E.C., Rolandsson, O., Hallmans, G., et al., 2009. Replication and extension of genome-wide association study results for obesity in 4923 adults from northern Sweden. Human Molecular Genetics 18(8): 1489–96, Doi: 10.1093/hmg/ddp041.

[45] Dempfle, A., Hinney, A., Heinzel-Gutenbrunner, M., Raab, M., Geller, F., Gudermann, T., et al., 2004. Large quantitative effect of melanocortin-4 receptor gene mutations on body mass index. Journal of Medical Genetics 41(10): 795–800, Doi: 10.1136/jmg.2004.018614.

[46] Heid, I.M., Vollmert, C., Hinney, A., Döring, A., Geller, F., Löwel, H., et al., 2005. Association of the 103I MC4R allele with decreased body mass in 7937 participants of two population based surveys. Journal of Medical Genetics: e21, Doi: 10.1136/jmg.2004.027011.

[47] Hoekstra, H.E., Hirschmann, R.J., Bundey, R.A., Insel, P.A., Crossland, J.P., 2006. A Single Amino Acid Mutation Contributes to Adaptive Beach Mouse Color Pattern. Science 313(5783): 101–4, Doi: 10.1126/science.1126121.

[48] Gonçalves, G.L., Paixão-Côrtes, V.R., Freitas, T.R.O., 2013. Molecular evolution of the melanocortin 1-receptor pigmentation gene in rodents. Genetics and Molecular Research 12(3): 3230–45, Doi: 10.4238/2013.February.28.24.

[49] Nachman, M.W., Hoekstra, H.E., D’Agostino, S.L., 2003. The genetic basis of adaptive melanism in pocket mice. Proceedings of the National Academy of Sciences of the United States of America 100(9): 5268–73, Doi: 10.1073/pnas.0431157100.

[50] Payne, J.L., Wagner, A., 2019. The causes of evolvability and their evolution. Nature Reviews Genetics: 24–38, Doi: 10.1038/s41576-018-0069-z.

[51] Zimmet, P., Thomas, C.R., 2003. Genotype, obesity and cardiovascular disease - Has technical and social advancement outstripped evolution? Journal of Internal Medicine: 114–25, Doi: 10.1046/j.1365-2796.2003.01170.x.

