## Supplementary figures and images for "Mouse models for V103I and I251L gain of function variants of the human MC4R display reduced adiposity and are not protected from a hypercaloric diet"

### Suppl. Figure 1

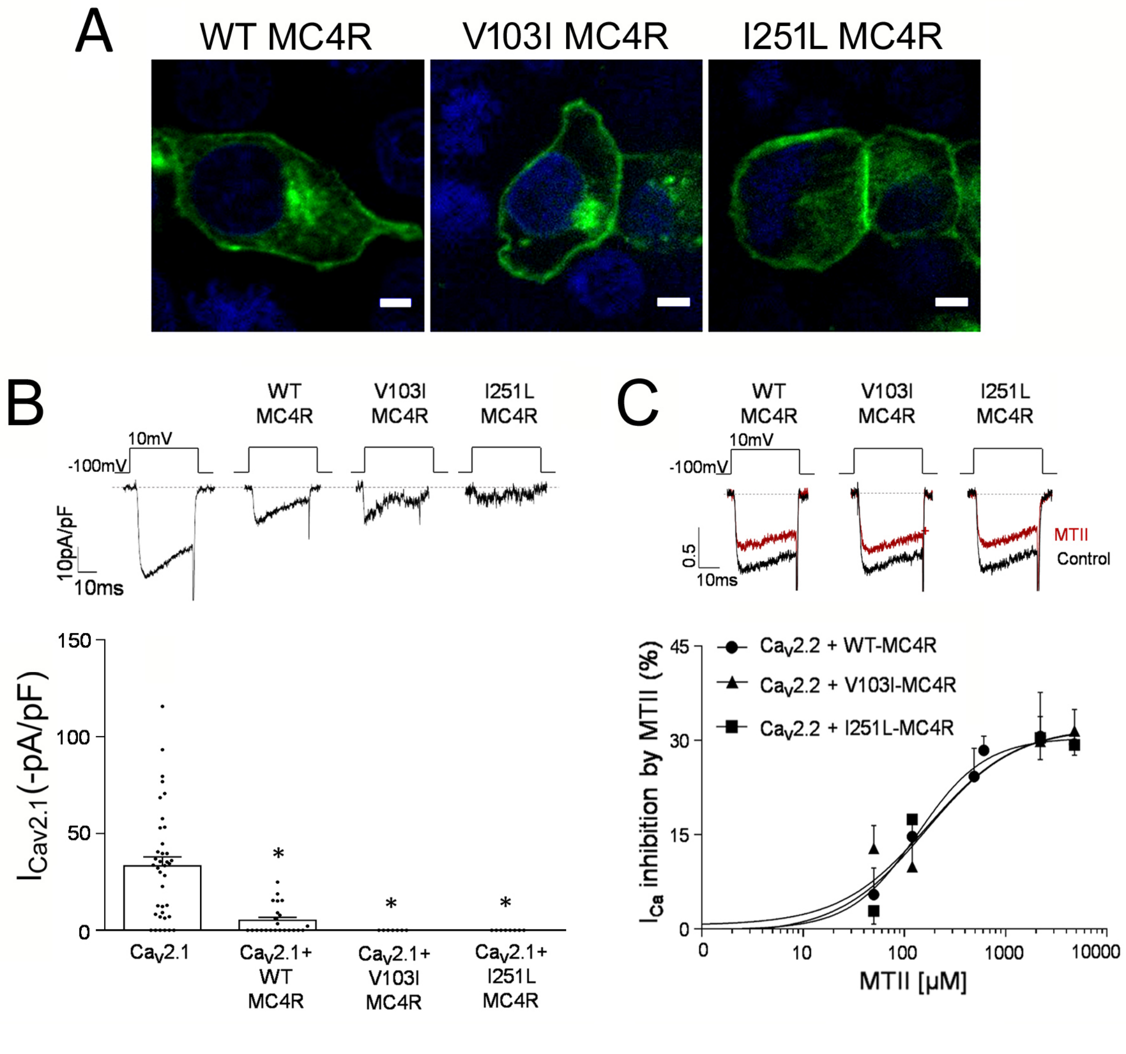


**Suppl. Figure S1**
