## Supplementary material for "Mouse models for V103I and I251L gain of function variants of the human MC4R display reduced adiposity and are not protected from a hypercaloric diet": Suppl. Table 1

**Table S1. ssODNs and primer sequences used to construct expression plasmids, sgRNAs, genotyping and qRT-PCRs.**

**ssODN *Mc4rV103I:***

5´-GTGAAGCTCTGGGCATCCGTATCCGTACTGTTTAACAGAGTAATGATGATGGTTTCCGACCCATTCGAAACGCTCACCAGCATATCTGCCACA-3´

**ssODN *Mc4rI251L:***

5´-AGTAAATGGAGAAAGAACGGGGCCCAGCAGACAACAAAGACTCCAAGCAGGATGGTCAATGTAATCGCCCCCTTCATGTTGGTACCCTGGCGG-3´

**Oligonucleotides for cloning the target specific crRNA region into DR274 plasmid:**

sgRNA *Mc4rV103I*

F: 5´-TAGGGCCGTATCCGTACTGTTTAAC-3´

R: 5´-AAACGTTAAACAGTACGGATACGGC-3´

sgRNA *Mc4rI251L*

F: 5´-TAGGGGACTCCAATCAGGATGGTCA-3´

R: 5´-AAACTGACCATCCTGATTGGAGTCC-3´

**Primer sequences for genotyping:**

***Mc4r-V103I***

F: 5´-AGCAACTTTTTGTTTCCCCCG-3´

R: 5´-GAGAGGCCATGAGAACTAGCA-3´

Size: +/+, +/V103I and V103I/V103I: 507 bp

F: 5´-ATGGGTCGGAAACCATCG-3´

R: 5´-CGAGTAAATGATGAAGAGGACGC-3´

Size: +/+ and +/V103I: 275 bp

F: 5´-ATGGGTCGGAAACCATCA-3´

R: 5´-CGAGTAAATGATGAAGAGGACGC-3´

Size: +/V103I and V103I/V103I: 275 bp

***Mc4r-I251L***

F: 5´-CCATTTGCAGCCTGCTTTCC-3´

R: 5´-CTTGACTCCGGAGGGCATAA-3´

Size: +/+, +/I251L and I251L/I251L: 516 bp

F: 5´-GATTACCTTGACCATCCTGA-3´

R: 5´-CTTGACTCCGGAGGGCATAA-3´

Size: +/+ and +/I251L: 191 bp

F: 5´-GATTACCTTGACCATCCTGC-3´

R: 5´-CTTGACTCCGGAGGGCATAA-3´

Size: +/I251L and I251L/I251L: 191 bp

**Primer sequences for qRT-PCR determinations:**

***Mc4r***

F: 5´-GTGATCGTGGCGATAGCCAA-3´

R: 5´-GGCATCCGTATCCGTACTGTT-3´

Size: 150 bp

***Actb***

F: 5’-AGAGGGAAATCGTGCGTGAC-3’

R: 5’-CAATAGTGATACCTGGCCGT-3’

Size: 138 bp

**Primers for expression plasmids construction:**

***Mc4r-V103I 1***

F: 5´-AAGCAGAGGAGGAGCCATTC -3´

R: 5´-TAACAGGGTAATGATGATGGTTTC-3´

Size: 842 bp

***Mc4r-V103I 2***

F: 5´-GAAACCATCATCATTACCCTGTTA-3´

R: 5´-GCATTACACAGAAGAGGCAGC-3´

Size: 803 bp

***Mc4r-V103I 3***

F: 5´-AAGCAGAGGAGGAGCCATTC -3´

R: 5´-GCATTACACAGAAGAGGCAGC-3´

Size: 1621 bp

***Mc4r-I251L 1***

F: 5´-AGCATTCAGAAGCACCAGCT-3´

R: 5´-CAAAGACTCCAAGCAGGATGGTCA-3´

Size: 992 bp

***Mc4r-I251L 2***

F: 5´-TGACCATCCTGCTTGGAGTCTTTG-3´

R: 5´-CTTAAAAAGAAGCATCAG-3´

Size: 820 bp

***Mc4r-I251L 3***

F: 5´-AGCATTCAGAAGCACCAGCT -3´

R: 5´-CTTAAAAAGAAGCATCAG -3´

Size: 1788 bp
